# NMDA receptor-targeted enrichment of CaMKIIα improves fear memory

**DOI:** 10.1101/2021.11.06.467164

**Authors:** Anthony Chifor, Jeongyoon Choi, Joongkyu Park

**Affiliations:** Department of Pharmacology, Wayne State University School of Medicine, Detroit, MI 48201, USA; Department of Neurology, Wayne State University School of Medicine, Detroit, MI 48201, USA

## Abstract

Calcium/calmodulin-dependent protein kinase II alpha (CaMKIIα) is an essential player in long-term potentiation and memory formation. However, the establishment of effective molecular interventions with CaMKIIα to improve memory remains a long-standing challenge. Here we report a novel intrabody targeting GluN1, a subunit of N-methyl-D-aspartate receptors (NMDARs). We identify this anti-GluN1 intrabody (termed <u>VHH A</u>nti-Glu<u>N1</u>; VHHAN1) by a synthetic phage display library selection and yeast-two-hybrid screenings. We validate specific targeting of VHHAN1 to GluN1 in heterologous cells and the mouse hippocampus. We further show that adeno-associated virus (AAV)-mediated expression of CaMKIIα fused with VHHAN1 is locally enriched at excitatory postsynaptic regions of the mouse hippocampus. We also find that the AAV- and VHHAN1-mediated postsynaptic enrichment of CaMKIIα in the hippocampus improves contextual fear memory in mice. This novel approach opens a new avenue to enhance memory ability in health and diseases.

## Introduction

Long-term potentiation (LTP) is an activity-dependent strengthening of synaptic transmission that has been considered a cellular mechanism of learning and memory^1,2^. In the hippocampus, LTP requires activation of N-methyl-D-aspartate receptors (NMDARs), a rise in postsynaptic calcium, and activation of calcium/calmodulin-dependent protein kinase II alpha (CaMKIIα)^1-6^. The importance of CaMKIIα in LTP and memory is well documented by knockout and inactive mutant (T286A) knock-in mouse studies^5-8^. Both knockout and mutant knock-in mice show a lack of hippocampal LTP and significant impairments in memory tests^5-8^. The critical links between CaMKIIα and LTP were further supported by recent works with CaMKIIα phosphorylation substrates (for example, TARPγ-8 and SynGAP) and a photoactivatable CaMKIIα inhibitor^9-11^.

However, little is known about how CaMKIIα can be utilized for memory improvements. This gap has been due to the limitations in our knowledge and techniques on the appropriate supply of exogenous CaMKIIα to facilitate molecular steps of memory formation. Virus-mediated hippocampal expression of wild-type CaMKIIα improves spatial memory in water maze tests of rats (∼1.15-fold longer time spent in the target area)^12^. Meanwhile, viral expression of the activated form of CaMKIIα (T286D/T305A/T306A) in the hippocampus impairs place avoidance memory and context discrimination in rodents similarly to what a dominant-negative form (K42M) caused^13,14^. We reason that the limited or no improvement in memory was due to the lack of (i) capability to respond to the postsynaptic calcium rise in the mutant forms and (ii) adequate postsynaptic supply of CaMKIIα as overexpressed CaMKIIα diffuses throughout neurons. Thus, we hypothesize that there is a need to provide (i) wild-type CaMKIIα with (ii) local postsynaptic enrichment proximal to NMDARs, where LTP initiates.

Here we report a novel VHH intrabody targeting the cytoplasmic region of a subunit of NMDARs GluN1 (termed <u>VHH A</u>nti-Glu<u>N1</u>; VHHAN1). We validated specific targeting of VHHAN1 to GluN1 in heterologous cells and the mouse hippocampus. Employing this intrabody, we supplied the adeno-associated virus (AAV)-mediated expression of CaMKIIα at excitatory postsynaptic regions of the mouse hippocampus. We further showed that this <u>C</u>aMKIIα <u>L</u>ocal <u>E</u>nrichment by <u>V</u>HH for <u>I</u>mprovement of memo<u>R</u>y (CLEVIR) in the hippocampus significantly enhances contextual memory in mice.

## Results

To develop an intrabody targeting endogenous NMDARs in the brain, it is critical to find an appropriate intracellular portion of NMDAR subunits for the intrabody binding. The heterotetrameric NMDARs consist of two subunits of GluN1 and two subunits of either GluN2 (GluN2A through GluN2D) or GluN3 (GluN3A and GluN3B)^15^. The cytoplasmic tail of GluN1 subunits provides a good platform for intrabody binding, but there are eight splice variants reported (Supplementary Fig. 1A). Therefore, as a target region, we chose a part of the mouse GluN1 cytoplasmic tail conserved throughout all the splice variants (C0 cassette, 834-863 amino acids of NP_032195.1) (Fig. 1A). We purified this fragment (His-AviTag-GluN1C0) from bacteria, immobilized it on streptavidin beads (Supplementary Figs. 1B,C), and conducted a nanobody screening with a synthetic phage display library (3.0 × 10^9^ nanobodies)^16^. We narrowed down nanobody candidates to 3.1 × 10^5^ by the first-phase phage display screening (Supplementary Fig. 1D). We further screened by yeast two-hybrid (Y2H) assays to search proteins that bind to the GluN1C0 intracellularly. We ran two rounds of Y2H screening with the C0 target region and the full-length cytoplasmic tail of GluN1 (834-938 amino acids of NP_032195.1). We obtained 118 positive clones from the Y2H screenings and confirmed three candidate clones after eliminating redundant clones by Sanger sequencing (Supplementary Fig. 1D).

**Fig. 1.**
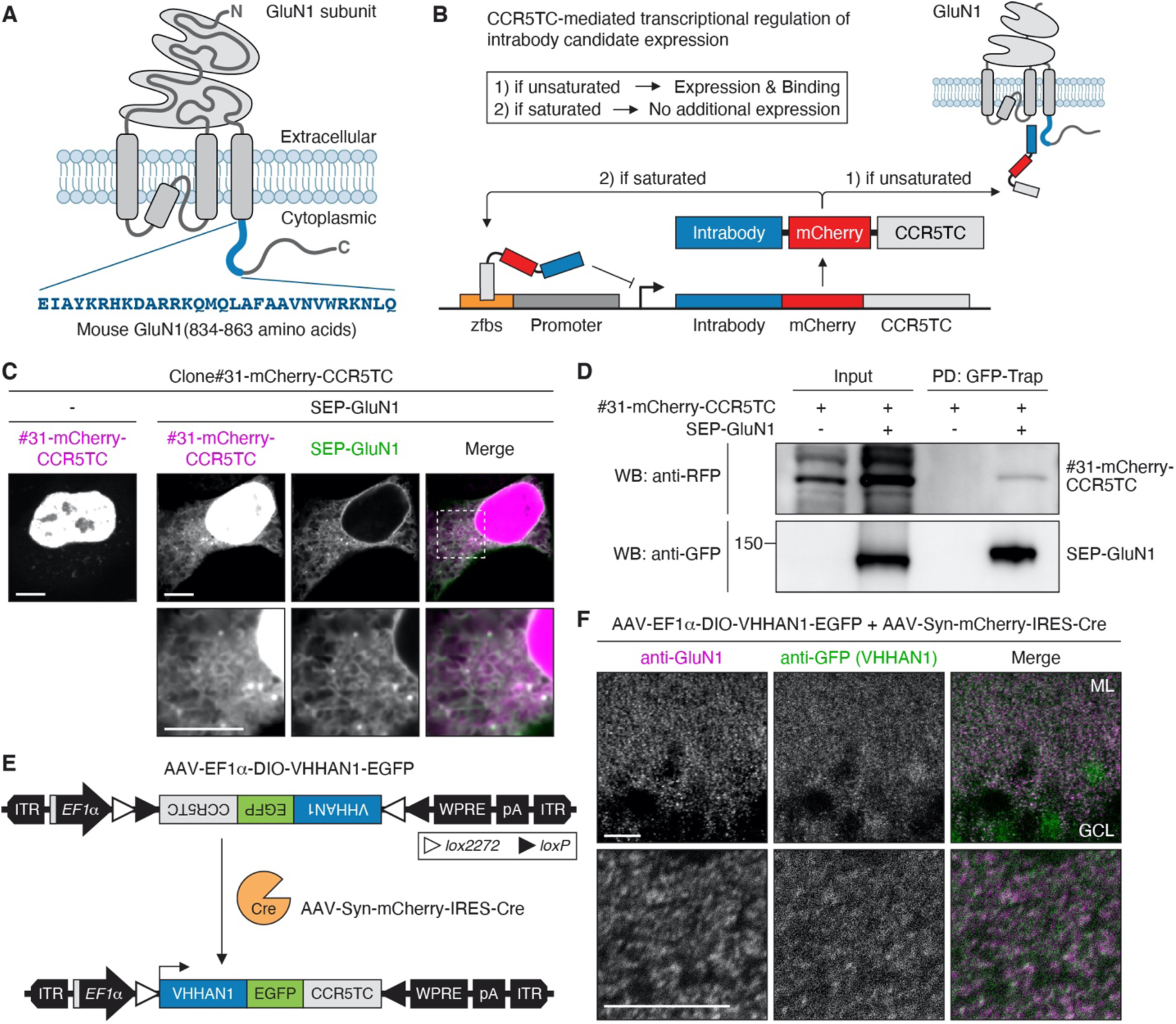
Development of anti-GluN1 intrabody. (**A**) Schematic diagram illustrating GluN1 domain structure and the anti-GluN1 intrabody target region (834-863 amino acids). (**B**) Illustration of CCR5TC-mediated transcriptional regulation. To avoid overproduction and subsequent random diffusion of intrabodies, intrabody candidates were fused with mCherry and CCR5TC. If the interaction is unsaturated, protein expression continues for further GluN1 binding. If 100% of GluN1 binds to the intrabody, then the newly synthesized intrabodies move to the nucleus and bind to a zinc-finger binding site (zfbs) upstream of a promoter to discontinue protein expression. (**C** and **D**) Colocalization and interaction of Clone#31 with GluN1 in heterologous cells. An intrabody candidate (Clone#31) was fused with mCherry and CCR5TC and expressed in HeLa cells without or with SEP-fused GluN1. In the absence of SEP-GluN1, Clone#31 localized in the nucleus due to the nuclear localization signal of CCR5TC (**C**, left panel). When co-expressed with SEP-GluN1 (green), Clone#31 (magenta) colocalized with SEP-GluN1 depicting the ER structure (**C**, right panel). GFP-Trap pull-down (PD) revealed the binding of Clone#31-mCherry-CCR5TC to SEP-GluN1 (**D**). (**E** and **F**) Colocalization of VHHAN1 (Clone#31) with endogenous GluN1 in the mouse hippocampus. Illustration of the AAV expressing EGFP-fused VHHAN1 (**E**). AAV-EF1α-DIO-VHHAN1-EGFP was co-injected with AAV-Syn-mCherry-IRES-Cre into the hippocampus to express VHHAN1-EGFP in hippocampal neurons (**E**). Anti-GFP immunostaining visualized postsynaptic puncta of VHHAN1 (green) that were colocalized with endogenous GluN1 puncta (magenta) in the dentate gyrus (**F**). ML, molecular layer; GCL, granule cell layer. Scale bar, 10 μm.

We next examined colocalization and interaction between GluN1 and the three intrabody candidates in a heterologous system. To circumvent the overproduction of intrabodies that results in diffusion throughout cells, we adopted a previously established system of CCR5TC transcriptional repression^17,18^. When an intrabody candidate expresses and binds to less than 100% of GluN1 (if unsaturated), protein expression continues for further GluN1 binding (Fig. 1B). When the interaction between GluN1 and the intrabody reaches 100% (if saturated), a newly expressed population of the intrabody candidate moves to the nucleus and binds to a zinc-finger binding site (zfbs) upstream of the promoter and stop further transcription (Fig. 1B). Using this system, we tested colocalization of super-ecliptic pHluorin (SEP)-fused GluN1^19^ and mCherry- and CCR5TC-fused intrabody candidates in HeLa cells. Spinning-disc confocal imaging revealed that a single expression of each intrabody candidate shows its enrichment in the nucleus due to the nuclear localization signal of CCR5TC (Fig. 1C and Supplementary Figs. 1E,F, left panels). However, co-expression of SEP-GluN1 in the endoplasmic reticulum (ER) recruited one of the candidates (Clone#31) effectively to show robust colocalization (Fig. 1C, right panel). The ER retention of GluN1 is expected in the absence of GluN2 subunits^20,21^. In contrast, the other two candidates failed or minimally colocalized with GluN1 (Supplementary Figs. 1E,F, right panels). Consistently, the interaction between SEP-GluN1 and Clone#31-mCherry-CCR5TC in HeLa cells was confirmed by a binding assay with GFP-Trap pull-down (Fig. 1D). The other intrabody candidates showed no or minimal interaction with GluN1 consistently with the colocalization data (Supplementary Figs. 1E-H). These data suggest that one of the intrabody candidates (Clone#31) colocalizes well and binds to GluN1 in heterologous cells, and we term this intrabody VHHAN1.

Finally, we validated postsynaptic targeting of VHHAN1 to endogenous GluN1 *in vivo*. To express VHHAN1 *in vivo*, we generated an AAV expressing enhanced green fluorescent protein (EGFP)-fused VHHAN1 (Fig. 1E). This AAV expresses VHHAN1-EGFP under an *elongation factor-1 alpha* (*EF1α*) promoter in a Cre recombinase (Cre)-dependent manner (Fig. 1E). To express VHHAN1 in neurons, we co-injected it with an AAV expressing Cre and mCherry under a neuronal promoter (*Synapsin*) (AAV-Syn-mCherry-IRES-Cre) (Fig. 1E). As shown in Fig. 1F, the mouse hippocampus expressing VHHAN1 showed robust colocalization of VHHAN1 with endogenous GluN1 as excitatory postsynaptic puncta. Taken together, we report the development of an intrabody VHHAN1 that targets endogenous GluN1 *in vivo*.

Using this intrabody, we tested whether wild-type CaMKIIα is locally enriched at excitatory postsynaptic regions by fusing with VHHAN1. We generated Cre-dependent AAVs expressing either HA-tagged CaMKIIα (AAV-Syn-DIO-HA-CaMKIIα) or both VHHAN1- and HA-tagged CaMKIIα under a *Synapsin* promoter (AAV-Syn-DIO-VHHAN1-HA-CaMKIIα) (Fig. 2A). The N-terminal modification of CaMKIIα with VHHAN1 and HA-tag was chosen to avoid potential disruption of the C-terminal dimerization of CaMKIIα hexamers^4^, and the N-terminal protein fusion is known to preserve the kinase activity^9^. Co-injections of these AAVs with AAV-Syn-mCherry-IRES-Cre allow us to achieve desired CaMKIIα expression and immediately visualize infected brain regions under a fluorescent microscope. As shown in Fig. 2B, the coverage of AAV infections in each hippocampus was symmetric within animals and comparable between animals. Immunohistochemistry with anti-HA antibody revealed that HA-CaMKIIα distributes intracellularly broader than VHHAN1-HA-CaMKIIα in infected neurons (Fig. 2C). Whereas the expression of HA-CaMKIIα showed robust signals in the medial prefrontal cortex (Fig. 2D, left panel) and the inner molecular layer of the hippocampal dentate gyrus (Fig. 2E, left panel), VHHAN1-HA-CaMKIIα did not show those patterns (Figs. 2D,E, right panels), indicating the widespread distribution of HA-CaMKIIα including axonal projections^22^. However, mCherry distribution in both conditions was even and comparable throughout the infected hippocampi (Fig. 2B), suggesting that the differences in anti-HA immunoreactivity in Figs. 2D,E result from the distinct subcellular distribution of VHHAN1-HA-CaMKIIα, not the AAV infection coverage. Focusing on the dentate gyrus, laser scanning confocal microscopy revealed that VHHAN1-HA-CaMKIIα shows regular punctate patterns in the molecular layers while HA-CaMKIIα displays diffused distribution following neurite projections (Fig. 2F). The puncta of VHHAN1-HA-CaMKIIα were colocalized well with an endogenous excitatory postsynaptic marker PSD-95 (Fig. 2F). These data suggest that VHHAN1 orients exogenous wild-type CaMKIIα to excitatory postsynaptic regions *in vivo*.

**Fig. 2.**
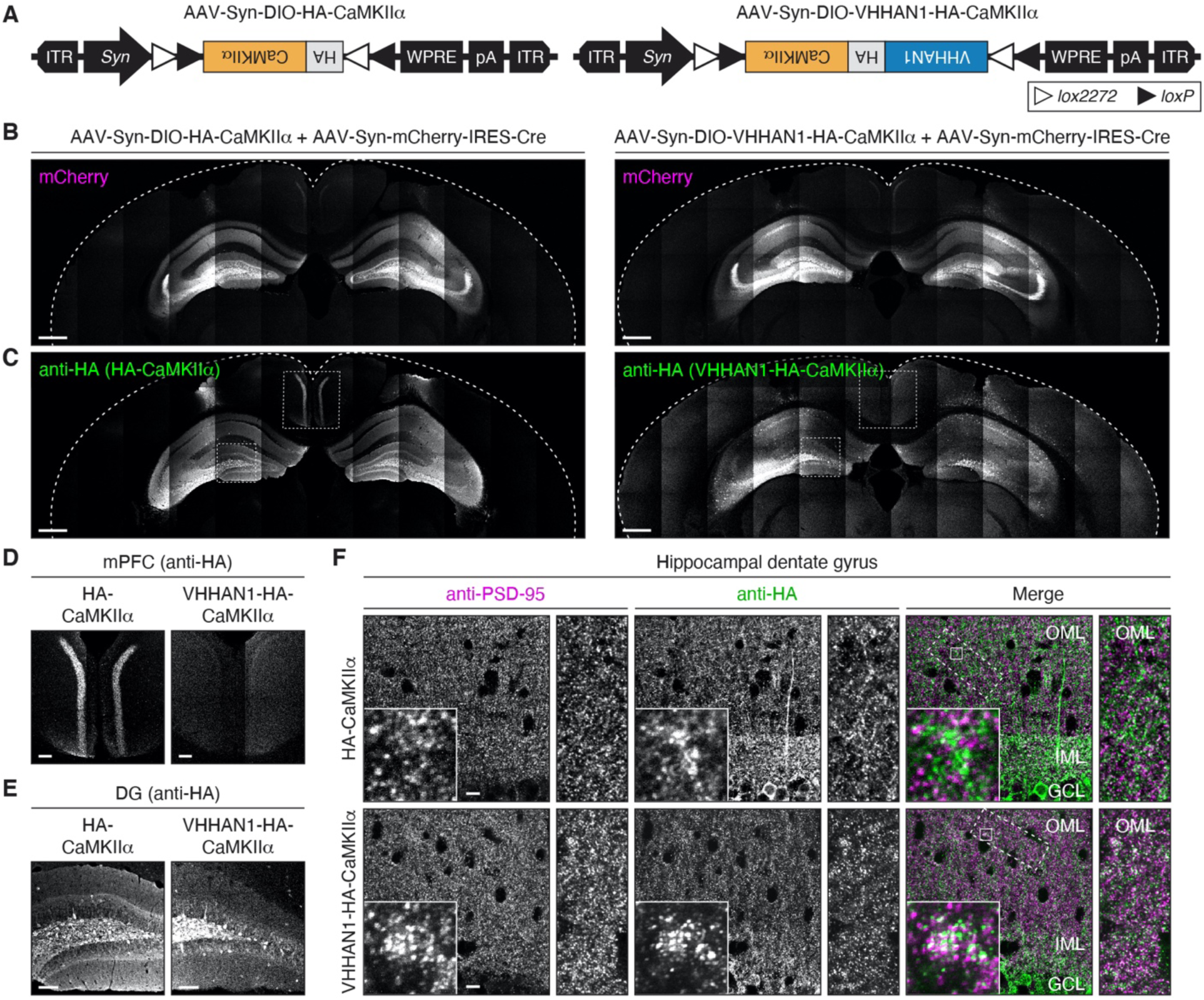
CaMKIIα local enrichment at excitatory postsynaptic regions by VHHAN1 in the mouse hippocampus. (**A**) Illustration of AAVs expressing HA-tagged CaMKIIα (left) and VHHAN1-HA-fused CaMKIIα (right). (**B**-**E**) Each AAV was co-injected with AAV-Syn-mCherry-IRES-Cre into the hippocampus to express HA-CaMKIIα (left) or VHHAN1-HA-CaMKIIα (right) in hippocampal neurons. The symmetric and comparable coverage of AAV infections in each hippocampus was validated by mCherry fluorescence (**B**). Anti-HA immunostaining visualized expression and localization of HA-CaMKIIα (left) and VHHAN1-HA-CaMKIIα (right) (**C**). The medial prefrontal cortex (mPFC) and hippocampal dentate gyrus (DG) regions are shown in (**D** and **E**) as indicated in dashed boxes in (**C**). Indicate the lack of presynaptic distribution of anti-HA signals in the mPFC and the inner molecular layer of DG from the VHHAN1-HA-CaMKIIα condition (**D** and **E**, right panels), indicating the postsynapse-oriented distribution of VHHAN1-HA-CaMKIIα. (**F**) High magnification confocal images of hippocampal DG co-stained with anti-HA and anti-PSD-95 (an excitatory postsynaptic marker) antibodies showed more regular punctate patterns of VHHAN1-HA-CaMKIIα, whereas HA-CaMKIIα displays more diffused distribution following neurite projections. Dashed and solid line boxes indicate the magnified regions for inset images. OML, outer molecular layer; IML, inner molecular layer; GCL, granule cell layer. Scale bar, 500 μm (**B** and **C**), 100 μm (**D** and **E**), and 10 μm (**F**).

Finally, we tested whether the VHHAN1-mediated local postsynaptic enrichment of wild-type CaMKIIα improves memory ability in mice using a fear conditioning paradigm. We co-injected Cre-dependent AAVs of either HA-CaMKIIα or VHHAN1-HA-CaMKIIα with AAV-Syn-mCherry-IRES-Cre into both hippocampi of wild-type mice as confirmed in Figs. 2B,C. After two weeks for recovery and protein expression, mice underwent a fear conditioning paradigm, then the next day, they received contextual and cued memory tests (Fig. 3A). Non-injected wild-type mice were tested as a control group. Overexpression of both HA-CaMKIIα and VHHAN1-HA-CaMKIIα had no significant impact on general locomotion compared to non-injected mice (Supplementary Fig. 2). Compared to HA-CaMKIIα and non-injected controls, the VHHAN1-HA-CaMKIIα group showed a significant increase in contextual memory (hippocampus-involved) (2.2-fold compared to HA-CaMKIIα and non-injected mice; *P* < 0.05) (Fig. 3B). In contrast, cued memory (amygdala-involved) was not significantly changed (Fig. 3C). The symmetric and comparable coverage of AAV infections in each mouse was confirmed by mCherry expression under a fluorescent microscope. Taken together, these data suggest that local postsynaptic enrichment of wild-type CaMKIIα proximal to NMDARs, where LTP occurs, enhances memory ability in mice. We term this molecular approach <u>C</u>aMKIIα <u>L</u>ocal <u>E</u>nrichment by <u>V</u>HH for <u>I</u>mprovement of memo<u>R</u>y (CLEVIR).

**Fig. 3.**
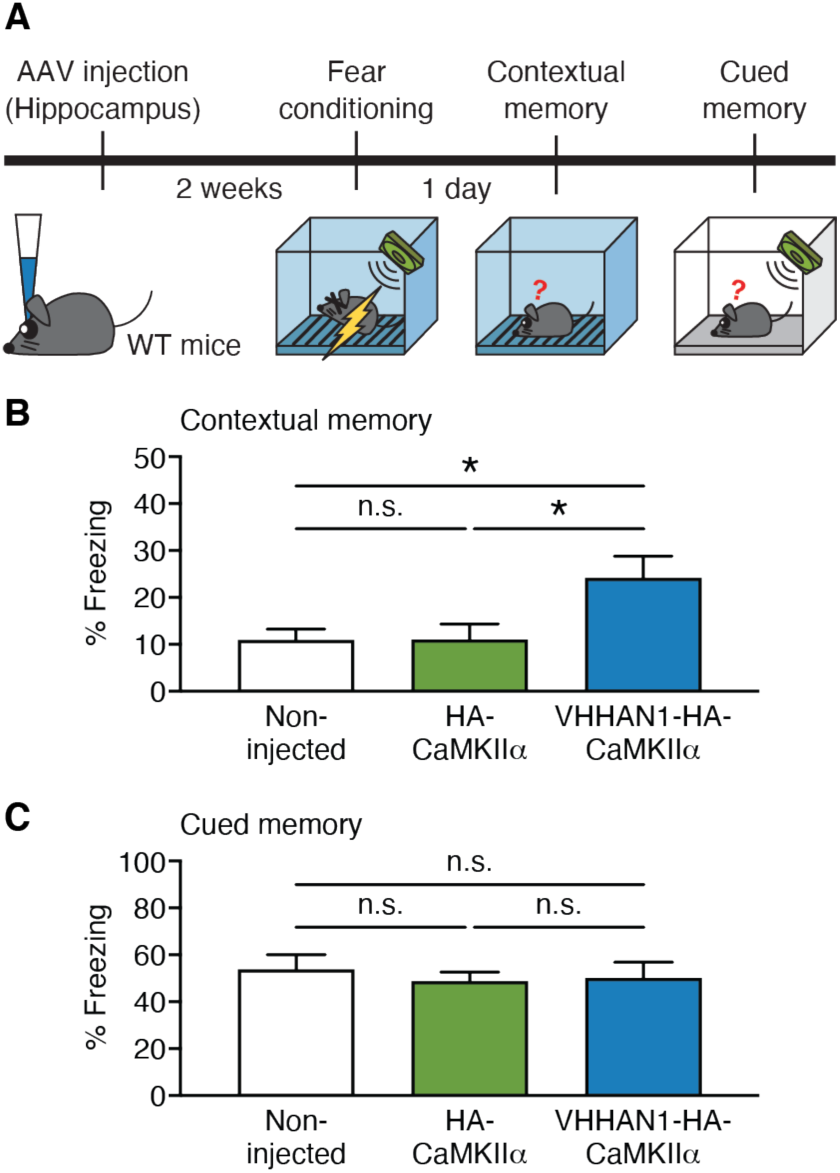
Improvement of contextual memory by VHHAN1-mediated postsynaptic enrichment of CaMKIIα in the hippocampus. (**A**) Schematic diagram depicting the experimental design. Either AAV-Syn-DIO-HA-CaMKIIα or AAV-Syn-DIO-VHHAN1-HA-CaMKIIα was co-injected with AAV-Syn-mCherry-IRES-Cre into the hippocampus of wild-type mice. After two weeks for protein expression, mice underwent a fear conditioning and testing paradigm. (**B**) Mice expressing VHHAN1-HA-CaMKIIα in the hippocampus showed an increase in contextual memory compared to non-injected and HA-CaMKIIα-expressing mice (*n* = 10 mice per group). (**C**) Cued fear memory showed no changes. Data are shown as mean ± SEM; \**P* < 0.05; n.s., not significant; Kruskal-Wallis tests with Dunn’s multiple comparisons.

## Discussion

Specific visualization and targeting of endogenous synaptic proteins in the brain are important to study synapse biology, but available molecular tools, including intrabodies, remain very limited. Our study reports a newly developed intrabody against the GluN1 subunit of NMDARs (i.e., VHHAN1). We demonstrated that this genetically encoded molecule could be readily fused with fluorescent proteins (mCherry and EGFP; Figs. 1C-F) and a synaptic enzyme (CaMKIIα; Fig. 2). Its small size (130 amino acids) is also beneficial to be delivered by AAVs, which have a limit in the length of the viral genome for packaging^23^. We validated the AAV-mediated visualization of endogenous GluN1 in the mouse hippocampus (Fig. 1F). We also showed that VHHAN1 drives the AAV-expressed CaMKIIα oriented towards excitatory postsynaptic regions in the mouse hippocampus while CaMKIIα alone diffuses throughout the infected neurons, including axonal projections (Fig. 2). Prior work developed two intrabodies against excitatory and inhibitory scaffolding proteins (PSD95.FingR and GPHN.FingR, respectively)^17,18^. Considering the differences in protein expression level, sub-synaptic localization, and functional roles of GluN1 and PSD-95^24-26^, it is plausible that each intrabody may be useful to investigate their sub-synaptic compartments in living neurons and animals.

Previous work showed a significant increase in spatial memory of rats by overexpression of wild-type CaMKIIα^12^. However, we observed no significant changes in contextual and cued fear memories with overexpression of HA-CaMKIIα alone (Figs. 3B,C). Capitalizing on VHHAN1, we supplied an exogenous population of wild-type CaMKIIα biased towards endogenous NMDARs, and the VHHAN1-mediated local enrichment of CaMKIIα (i.e., CLEVIR) in the hippocampus significantly improved contextual memory more than 2-fold (Figs. 2,3). This finding implies that there may be negative impacts from non-synaptic overexpression of CaMKIIα on encoding memories.

Collectively, these molecular tools (VHHAN1 and CLEVIR) are likely to provide a wide range of experimental potential to investigate synapse and memory biology in the field.

## Acknowledgements

We thank Drs. Rodrigo Andrade, Michael J. Bannon, and Sokol V. Todi for providing valuable input. We also thank members of Park lab for helpful discussions. This work was supported by NIH R21AG068423 (J.P.).

## Author contributions

A.C. and J.P. initiated and conceived the study. A.C., J.C, and J.P. designed and performed the experiments. A.C., J.C., and J.P. analyzed the data. J.P. wrote the manuscript. All authors discussed the results and contributed to the manuscript.

## Competing interests

The authors declare no competing interests.

---

**Supplementary Fig. 1 (Related to Fig. 1).**
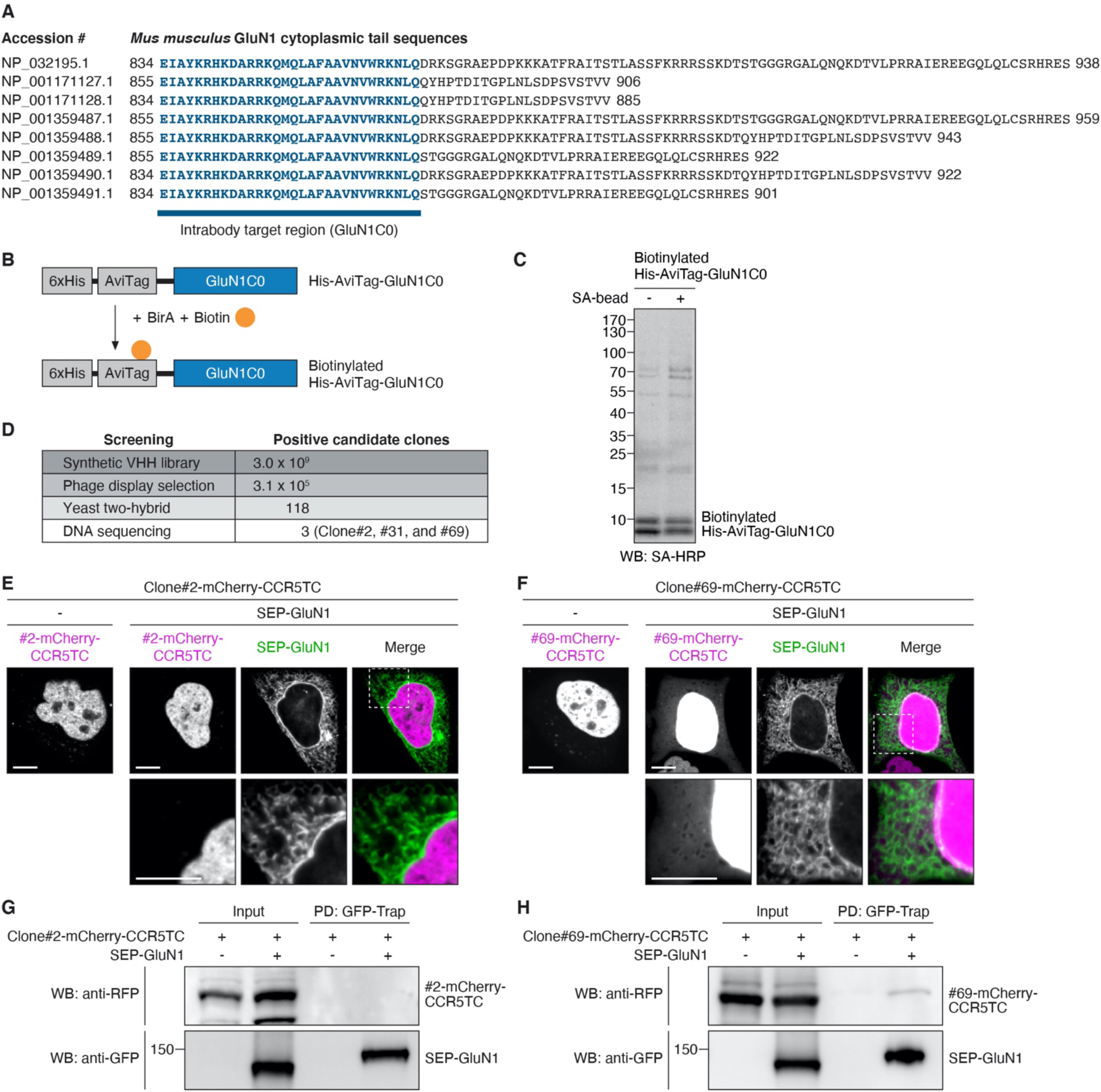
Validation of anti-GluN1 intrabody candidates. (**A**) Comparison of eight splice variants of mouse GluN1 cytoplasmic tail sequences. The conserved region (834-863 amino acids of NP_032195.1; C0 cassette) was chosen as the intrabody target (indicated in blue). (**B** and **C**) The bacterially purified His-AviTag-tagged GluN1C0 fragment (∼11 kDa) was biotinylated by a BirA enzyme *in vitro* (**B**) and immobilized on streptavidin beads (SA-bead) (**C**). *In vitro* biotinylation of His-AviTag-GluN1C0 fragment (**C**, left lane) and immobilization on SA-beads (**C**, right lane) were confirmed by Western blot using streptavidin-horseradish peroxidase conjugates (SA-HRP). (**D**) Summary of positive clone numbers of anti-GluN1 intrabody candidates after each screening. (**E** and **F**) Colocalization tests of Clone#2 and #69 with GluN1 in heterologous cells. Single transfection of DNA constructs encoding mCherry- and CCR5TC-fused Clone#2 or #69 in HeLa cells showed their nuclear localization due to the CCR5TC (**E** and **F**, left panels). Co-expression of SEP-GluN1 (green) showed no or minimal colocalization of the intrabody candidates (magenta; **E** and **F**, right panels). (**G** and **H**) Binding tests of Clone#2 and #69 with GluN1 in heterologous cells. GFP-Trap pull-down (PD) revealed that Clone#2-mCherry-CCR5TC does not bind to SEP-GluN1 in HeLa cells (**G**) consistently with the colocalization test data (**E**). GFP-Trap PD showed that Clone#69-mCherry-CCR5TC can bind to SEP-GluN1 in HeLa cells (**H**), but Clone#69 was not selected as the anti-GluN1 intrabody because it did not show robust colocalization with SEP-GluN1 (**F**). Scale bar, 10 μm.

**Supplementary Fig. 2 (Related to Fig. 3).**
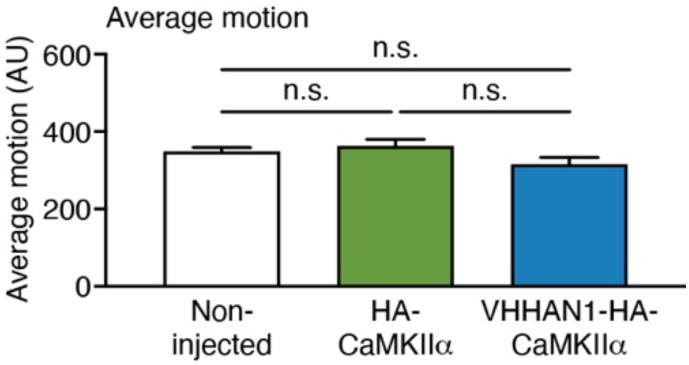
General locomotor activity. There were no significant differences in general locomotor activity (average motion; arbitrary unit) measured during the habituation session (after two-week protein expression). Data are shown as mean ± SEM; *n* = 10 mice per group; n.s., not significant; Kruskal-Wallis tests with Dunn’s multiple comparisons.

## Methods

### Plasmids

To express 6xHis- and AviTag-tagged mouse GluN1 C0 cassette fragment (834-863 amino acids; NP_032195.1) in bacteria, a DNA fragment encoding AviTag-GluN1(834-863) was synthesized by the Integrated DNA Technologies (Table 3) and inserted into pET-28a vector (Novagen) using NdeI and XhoI sites.

For mammalian expression of anti-GluN1 intrabody candidates, a CCR5TC transcriptional repression system (a zinc-finger DNA-binding domain and a KRAB(A) transcriptional repressor) was adopted^17^. Intermediate constructs were generated by inserting each intrabody candidate (Clone#2, #31, and #69) and an mCherry fragment into a linearized pcDNA3 vector using HindIII and XbaI sites. A flexible linker of Gly-Ser-Gly-Ser-Gly was inserted between the intrabody candidates and mCherry. Then, the fragment of intrabody candidate-mCherry was connected with the CCR5TC fragment by overlap extension polymerase chain reaction (PCR) with primers listed in Table 2. The amplified fragments were inserted using KpnI and MluI sites into the pCAG vector backbone containing a zinc-finger binding site upstream of a *CAG* promoter prepared from pCAG-GPHN.FingR-EGFP-CCR5TC (Addgene #46296)^17^.

**Table 1.** Reagent list and suppliers.

| Reagent or Resource | Source | Identifier |
| --- | --- | --- |
| **Antibodies** |  |  |
| Mouse monoclonal anti-RFP (clone 6G6) | Chromotek | Cat# 6G6 |
| Rabbit polyclonal anti-GFP | SYSY | Cat# 132003 |
| Mouse monoclonal anti-NMDAR1 (clone 54.1) | Millipore | Cat# MAB363 |
| Rat monoclonal anti-HA (clone 3F10) | Roche | Cat# ROAHAHA |
| Mouse monoclonal anti-HA (clone 16B12) | BioLegend | Cat# 901514 |
| Guinea pig polyclonal anti-PSD-95 | Frontier Institute | Cat# PSD95-GP-Af660 |
| Goat anti-rat IgG, Alexa Fluor 488 | ThermoFisher Scientific | Cat# A21208 |
| Goat anti-rabbit IgG, Alexa Fluor 594 | ThermoFisher Scientific | Cat# A11037 |
| Goat anti-mouse IgG, Alexa Fluor 647 | ThermoFisher Scientific | Cat# A32728 |
| Goat anti-guinea pig IgG, Alexa Fluor 647 | ThermoFisher Scientific | Cat# A21450 |
| Peroxidase AffiniPure goat anti-mouse IgG | Jackson ImmunoResearch | Cat# 115-035-146 |
| Peroxidase AffiniPure goat anti-rabbit IgG | Jackson ImmunoResearch | Cat# 111-035-144 |
| **Virus Strains** |  |  |
| AAV-EF1α-DIO-VHHAN1-EGFP | This paper | N/A |
| AAV-Syn-DIO-HA-CaMKIIα | This paper | N/A |
| AAV-Syn-DIO-VHHAN1-HA-CaMKIIα | This paper | N/A |
| AAV-Syn-mCherry-IRES-Cre | This paper | N/A |
| **Chemicals** |  |  |
| Ni-NTA agarose | Qiagen | Cat# 30210 |
| Gateway™ LR Clonase™ II Enzyme mix | Invitrogen | Cat# 11791020 |
| GFP-Trap magnetic agarose | Chromotek | Cat# gtma-20 |
| Anti-HA magnetic beads | ThermoFisher Scientific | Cat# 88837 |
| FuGENE HD transfection reagent | Promega | Cat# E2311 |
| Benzonase nuclease | Sigma-Aldrich | Cat# E1014 |
| SuperSignal™ West Pico PLUS Chemiluminescent Substrate | ThermoFisher Scientific | Cat# 34580 |
| Paraformaldehyde | Sigma-Aldrich | Cat# P6148 |
| Pepsin | Agilent | Cat# S300230-2 |
| Normal goat serum blocking solution | Vector Laboratories | Cat# S-1000 |
| DAPI Fluoromount-G | SouthernBiotech | Cat# 0100-20 |
| HiTrap heparin high performance columns | GE Healthcare | Cat# 17-0406-01 |
| **Experimental Models: Cell Lines** |  |  |
| HeLa cells | ATCC | Cat# CCL-2 |
| 293FT cell line | Invitrogen | Cat# R70007 |
| BL21-CodonPlus (DE3)-RIPL competent cells | Agilent | Cat# 230280 |

| Experimental Models: Organisms/Strains |  |  |
| --- | --- | --- |
| C57BL/6J mice | The Jackson Laboratory | Stock# 000664 |
| Oligonucleotides |  |  |
| Primers for DNA constructs, see Table 2 | This paper | N/A |
| Recombinant DNA |  |  |
| pET-28a-AviTag-GluN1(834-863) | This paper | N/A |
| pCAG-VHHAN1(Clone#31)-mCherry-CCR5TC | This paper | N/A |
| pCAG-Clone#2-mCherry-CCR5TC | This paper | N/A |
| pCAG-Clone#69-mCherry-CCR5TC | This paper | N/A |
| pAAV-EF1α-DIO-VHHAN1-EGFP | This paper | N/A |
| pAAV-Syn-DIO-HA-CaMKIIα | This paper | N/A |
| pAAV-Syn-DIO-VHHAN1-HA-CaMKIIα | This paper | N/A |
| pAAV-Syn-mCherry-IRES-Cre | This paper | N/A |
| pCI-SEP-NRI | Kopec et al., 2006 | RRID: Addgene_23999 |
| Software and Algorithms |  |  |
| Fiji | Schindelin et al., 2012 | http://fiji.sc |

**Table 2.**
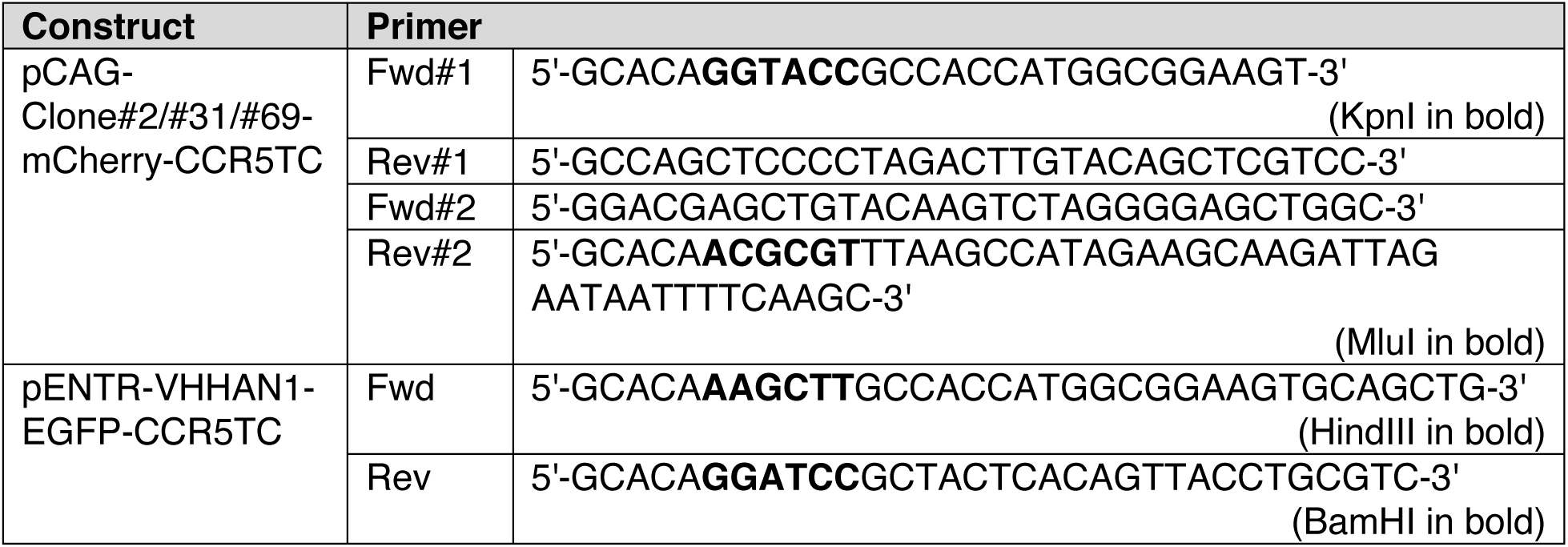
Primers used for plasmid construction.

To generate an AAV construct expressing VHHAN1-fused EGFP, we first generated a pENTR construct that contains VHHAN1 and EGFP-CCR5TC fragments. The VHHAN1 fragment was amplified by PCR with primers listed in Table 2 and digested with HindIII and BamHI. The EGFP-CCR5TC fragment was obtained by a BamHI and XhoI digestion from pCAG-GPHN.FingR-EGFP-CCR5TC (Addgene #46296)^17^. The DNA fragments of VHHAN1 and EGFP-CCR5TC were inserted into a pENTR vector using HindIII and XhoI sites. This entry clone was recombined by Gateway LR Clonase II enzyme (Invitrogen) with a pAAV-EF1α-DIO-Dest vector, which was generated by replacing a cassette of a ccdB gene and attR1 and attR2 sites with the coding region of AAV-EF1α-DIO-PSD95.FingR-EGFP-CCR5TC (Addgene #126216)^18^.

For the cistronic expression of mCherry and Cre under a *Synapsin* promoter, we first inserted a DNA fragment encoding mCherry-IRES-Cre (synthesized by the Integrated DNA Technologies; Table 3) into pENTR vector using BamHI and EcoRI sites. We then performed Gateway LR recombination of pENTR-mCherry-IRES-Cre and pAAV-Syn-Dest constructs.

**Table 3.**
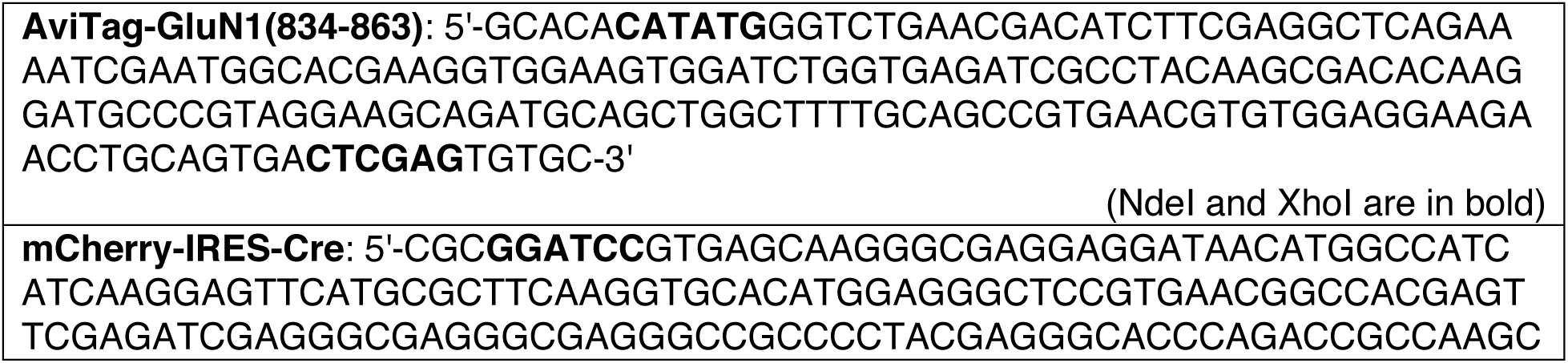
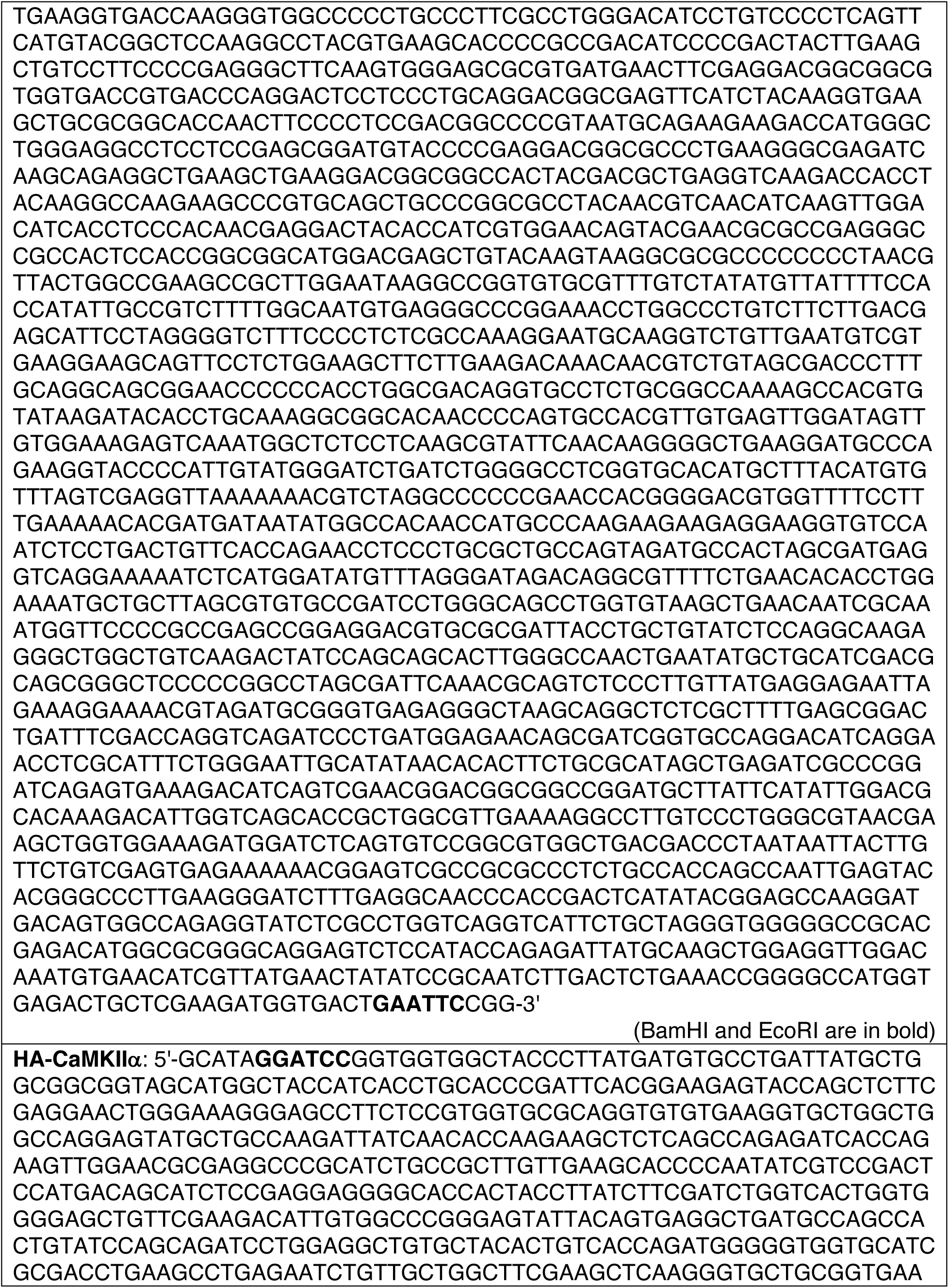
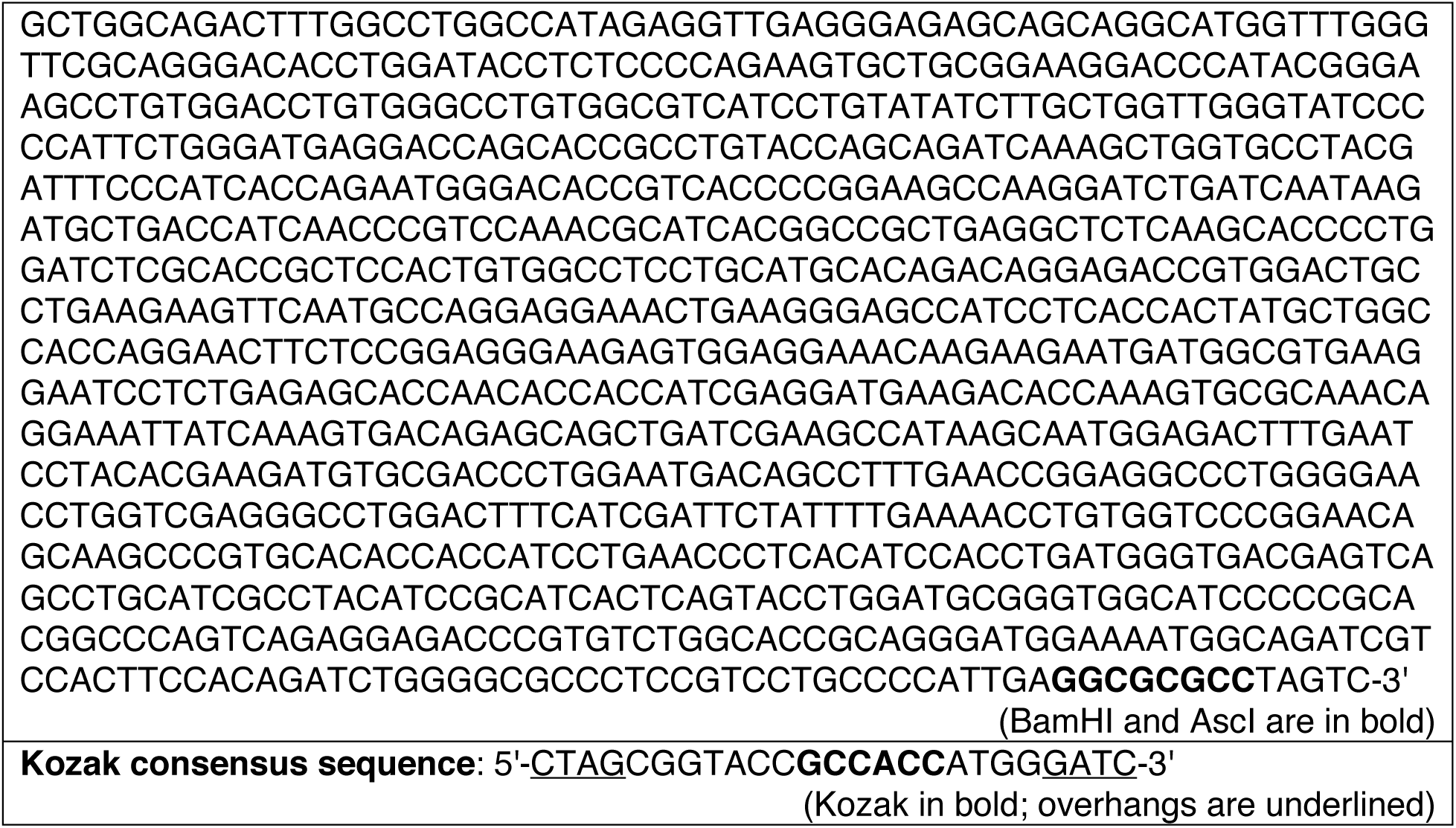
Synthetic DNAs used in this work.

For the expression of VHHAN1-HA-CaMKIIα under a *Synapsin* promoter, an intermediate construct was first generated by Gateway LR recombination of pENTR-VHHAN1 and pAAV-Syn-DIO-Dest constructs, and then a DNA fragment encoding HA-CaMKIIα (synthesized by the Integrated DNA Technologies; Table 3) was inserted next to VHHAN1 using BamHI and AscI sites. To express HA-CaMKIIα alone as a control, the DNA fragment encoding VHHAN1 was removed by a BamHI and NheI digestion and replaced with an annealed primer linker encoding the Kozak consensus sequence (Table 3).

The pCAG-GPHN.FingR-EGFP-CCR5TC, AAV-EF1α-DIO-PSD95.FingR-EGFP-CCR5TC, and pCI-SEP-NRI were gifts from Don Arnold (Addgene plasmid #46296; http://n2t.net/addgene:46296; RRID: Addgene_46296), Xue Han (Addgene plasmid #126216; http://n2t.net/addgene:126216; RRID: Addgene_126216), and Robert Malinow (Addgene plasmid #23999; http://n2t.net/addgene:23999; RRID: Addgene_23999), respectively.

### Preparation of anti-GluN1 intrabody target fragment

To design an anti-GluN1 intrabody target fragment, eight splice variants of the mouse GluN1 cytoplasmic tail that are deposited in the NCBI Reference Sequences database (NP_032195.1, NP_001171127.1, NP_001171128.1, NP_001359487.1, NP_001359488.1, NP_001359489.1, NP_001359490.1, and NP_001359491.1) were compared (Supplementary Fig. 1A). The conserved region (834-863 amino acids; NP_032195.1) was fused with a six histidine tag (6xHis) and an AviTag (pET-28a-AviTag-GluN1C0) and expressed in BL21-CodonPlus (DE3)-RIPL cells (Agilent). Expression of His-AviTag-GluN1(834-863) fragment was induced by 1 mM isopropyl b-D-thiogalactopyranoside at 37°C for 3.5 h, and the fragment was purified with Ni-NTA agarose (Qiagen). All eluted samples were then desalted using an Econo-Pac 10DG desalting column (Bio-Rad) and concentrated using an Amicon ultra-centrifugal filter unit (Millipore). The purified fragment was confirmed by resolving on a 17% SDS-PAGE gel and Coomassie Brilliant Blue G-250 staining.

### Phage display selection and yeast two-hybrid (Y2H) screening

The BirA-mediated *in vitro* biotinylation of the GluN1(834-863) fragment was performed by Hybrigenics Services SAS (www.hybrigenics-services.com). The first-phase screening was conducted using a hs2dAb synthetic library of humanized nanobodies (3.0 × 10^9^ nanobodies)^16^. For the Y2H screen, the coding sequences for the C0 cassette (834-863 amino acids; NP_032195.1) and the full-length cytoplasmic tail of mouse GluN1 (834-938 amino acids; NP_032195.1) were sub-cloned into pB27 as a C-terminal fusion to LexA. Two rounds of Y2H screenings were performed by Hybrigenics Services SAS.

### Cell culture and transfection

HeLa and 293FT cells were maintained in DMEM supplemented with 10% (v/v) fetal bovine serum, 100 U/mL penicillin-streptomycin, and 2 mM L-Glutamine in a humidified incubator with 37°C and 5% CO_2_. For confocal imaging and binding assays, transient transfection was performed using FuGENE HD transfection reagent according to the manufacturer’s instructions (Promega).

### Cell imaging

Cells were plated on 35-mm glass-bottom dishes (MatTek Corporation) and imaged under a spinning-disc confocal microscope (Zeiss Cell) equipped with a 63x (NA 1.4) objective approximately 18 h after transfection. Before imaging, cells were transferred to a pre-warmed buffer containing 10 mM HEPES (pH 7.4), 137 mM NaCl, 2.5 mM KCl, 2 mM CaCl_2_, and 1.3 mM MgCl_2_.

### Binding assay and Western blot

Each plasmid encoding an mCherry-CCR5TC-fused anti-GluN1 intrabody candidate (#2, #31, or #69; 2 μg) was co-transfected into HeLa cells (at ∼70% confluency in 35-mm dishes) with or without pCI-SEP-NRI (encoding SEP-GluN1; 1 μg) using FuGENE HD transfection reagent. Cells were lysed in 1 ml of RIPA buffer (50 mM Tris, pH 8.0, 150 mM NaCl, 1% Triton X-100, 0.5% sodium deoxycholate, and 0.2% SDS) containing ∼750 U Benzonase nuclease. After clearing up by centrifugation (15,000 *g* for 15 min at 4°C), each lysate was mixed with 15 μl of GFP-Trap magnetic beads overnight at 4°C. The protein complex pulled down by GFP-Trap beads (Chromotek) was washed three times with RIPA buffer and eluted in 30 μl of 1x Laemmli sample buffers by heating at 75°C for 10 min.

The eluted protein complexes were resolved on 7% SDS-PAGE gels and analyzed by Western blot. Briefly, SDS-PAGE gels were transferred to polyvinylidene difluoride membranes, and the membranes were blocked with 3% non-fat dry milk in TBST buffer (20 mM Tris, 150 mM NaCl, and 0.1% Tween 20) for 1 h at room temperature. Membranes were then probed overnight at 4°C with TBST buffer containing 1% non-fat dry milk and primary antibodies: mouse monoclonal anti-RFP (Chromotek, clone 6G6, 1:3,000), rabbit polyclonal anti-GFP (SYSY, 1:3,000), mouse monoclonal anti-NMDAR1 (Millipore, clone 54.1, 1:3,000), and mouse monoclonal anti-HA (BioLegend, clone 16B12, 1:3,000). The next day, membranes were washed three times in TBST buffer and incubated for 2 h at room temperature with TBST buffer containing 1% non-fat dry milk and horseradish peroxidase (HRP)-conjugated anti-mouse IgG or anti-rabbit IgG antibodies. The membranes were then washed three times with TBST buffer, and signals were visualized with an enhanced chemiluminescence reagent (Thermo Scientific, SuperSignal™ West Pico PLUS Chemiluminescent Substrate).

### Animals

C57BL/6J wild-type mice were obtained from the Jackson Laboratory (Stock# 000664). Mice were maintained at the Division of Laboratory Animal Resources facility of Wayne State University and treated under the guidelines of the Institutional Animal Care and Use Committee of Wayne State University.

### AAV production

Adeno-associated virus (AAV) was generated using the AAV-DJ Helper Free system (Cell Biolabs) as published previously^9,27^. Each AAV construct was co-transfected with pAAV-DJ and pHelper into 293FT cells (Invitrogen). Transfected cells were lysed three days after transfection, and AAVs were purified with 1-ml HiTrap heparin high-performance columns (GE Healthcare) as described^27^. The titer of each AAV was determined by semi-quantitative PCR.

### AAV injection

Stereotaxic injections of mouse hippocampi were performed as described previously^9^. The 6- to 9-week-old C57BL/6 wild-type mice were anesthetized with 1.5-3.0% isoflurane and placed in a stereotaxic apparatus (Kopf Instruments). The skull was exposed over the hippocampi based on stereotactic coordinates. Then, 0.7-1.0 μl of AAV was injected into the dorsal hippocampi using a glass pipette (tip diameter 7∼10 μm) at a rate of 100 nl/min using a syringe pump (Micro2T; World Precision Instruments). We injected each AAV with the following titer per hippocampus: 1.0 × 10^6^ viral genome (vg) of AAV-EF1α-DIO-VHHAN1-EGFP and 4.0 × 10^7^ vg of AAV-Syn-mCherry-IRES-Cre for Fig. 1F; 3.8 × 10^7^ vg of AAV-Syn-DIO-HA-CaMKIIα, 2.1 × 10^7^ vg of AAV-Syn-DIO-VHHAN1-HA-CaMKIIα, and 2.7 × 10^7^ vg of AAV-Syn-mCherry-IRES-Cre for Figs. 2,3. The injection site was standardized among animals by using stereotaxic coordinates (ML = ±2.00, AP = –2.20, DV = –1.90, –1.65, and –1.40 for hippocampi) from bregma. At the end of the injections, we waited 10 min before retracting the pipette. One to two weeks after injections, mice were subjected to tissue harvest or behavioral tests.

### Immunohistochemistry

Mice were deeply anesthetized and transcardially perfused with 4% paraformaldehyde (PFA) in 0.1 M phosphate buffer (PB, pH 7.4). After postfixation for either 3 h (for GluN1) or overnight (for other proteins), and 50 μm thick coronal sections were cut on a vibratome (Leica) at 4°C. To stain postsynaptic proteins, free-floating sections were treated with 1 mg/ml pepsin (Agilent) in 0.2 N HCl at 37°C for 2 min for antigen retrieval and washed in PBS three times. The slices were blocked for 45 min in PBS containing 5% normal goat serum (Vector Laboratories) and 0.3% Triton X-100 and incubated overnight at room temperature with PBS containing 3% normal goat serum, 0.1% Triton X-100, and primary antibodies: rabbit polyclonal anti-GFP (SYSY, 1:1,000), mouse monoclonal anti-NMDAR1 (Millipore, clone 54.1, 1:1,000), rat monoclonal anti-HA (Roche, clone 3F10, 1:1,000), and guinea pig polyclonal anti-PSD-95 (Frontier Institute, 1:1,000). Sections were then washed three times with PBS and incubated for 2 h at room temperature with secondary antibodies: Alexa Fluor 594-conjugated goat anti-rabbit IgG (H+L) highly cross-adsorbed secondary antibody, Alexa Fluor 647-conjugated goat anti-mouse IgG (H+L) highly cross-adsorbed secondary antibody, Alexa Fluor 488-conjugated donkey anti-rat IgG (H+L) highly cross-adsorbed secondary antibody, and Alexa Fluor 647-conjugated goat anti-guinea pig IgG (H+L) highly cross-adsorbed secondary antibody. Then, sections were washed three times in PBS and mounted on microscope slides with DAPI Fluoromount-G (SouthernBiotech). Confocal fluorescence images were acquired on a laser scanning confocal microscope (LSM 780; Zeiss) equipped with a 10x (NA 0.3) objective or a 63x (NA 1.4) objective. Images were edited and pseudo-colored using Fiji (http://fiji.sc).

### Behaviors

All mice were housed on a 12-h light-dark cycle with food and water ad libitum. All behavioral experiments were conducted with 8- to 11-week-old littermates at the time of testing and randomization of males and females. All behavioral tests and analyses were performed in a blind manner to the AAV genotypes. Mice were tested in four sessions (habituation, training, contextual memory test, and cued memory test) as described previously^9^. For habituation, mice were exposed to a dark plastic chamber with a plastic A-shaped frame insert and a plastic floor cover (Med associates) (as context A). No shocks or cues were delivered. The average motion values in this session were used to monitor changes in general locomotor activity by overexpression of HA-CaMKIIα or VHHAN1-HA-CaMKIIα. (Supplementary Fig. 2). In the training sessions, mice were placed in a modified plastic chamber with a house light, removal of the A-frame insert, and grids for shock delivery (as context B). After 160 sec as the baseline, mice underwent three repeats (with 40-sec intertrial intervals) of a 20-sec tone (85 dB, 2.8 kHz) and a 2-sec foot shock (0.5 mA) coincided at the end of each tone. After training, mice were returned to their home cage. The next day, mice were placed again in the same training chamber (context B) to test contextual memory for 6 min. To test cued memory, mice were placed in the context A chamber, and the three rounds of tones (20 sec, 85 dB, 2.8 kHz) and intertrial intervals (40 sec) were administered as described in the training sessions without foot shocks. Every session was filmed by an infrared video camera (Med associates), and freezing behavior (defined as complete lack of movement between every 1-sec frame of videos) was analyzed by Video Freeze (Med associates) software in a blind manner. No obvious differences in body weight and responses to shocks were observed. Brains from all tested mice were subjected to brain slice sectioning and fluorescent microscopy to confirm the symmetric AAV injections in both hippocampi.

### Statistical analysis

All data are given as mean ± SEM. The sample size was chosen based on the previous studies^9^. Statistical significance between means was calculated using Kruskal-Wallis tests with Dunn’s multiple comparisons. Statistical significance is indicated as follows: \**P* < 0.05 and n.s., not significant.

## Notes

### Competing Interest Statement

The authors have declared no competing interest.

